# Combining amino acid frequency and 1D convolutional neural network embeddings for the identification of protein-protein interactions using a random forest classifier

**DOI:** 10.64898/2026.05.15.725340

**Authors:** Nisa A. Sindhi, Nikhil M. Pawar, Jamie D. Dixson, Dana M. García

## Abstract

Predicting protein–protein interactions is a fundamental problem in molecular biology. Experimental approaches for identifying protein-protein interactions are time-consuming and labor-intensive, motivating the development of efficient computational alternatives, including machine learning-based methods. However, conventional machine learning methods often rely on manually engineered features that require substantial domain expertise. In this study, we propose a two-stage framework to address these limitations. In the first stage, a one-dimensional convolutional neural network autoencoder is used to automatically learn latent representations from protein sequences. The quality of these features is evaluated through reconstruction error, reflecting how accurately the model reconstructs the original sequence. In the second stage, these learned features are combined with amino acid frequency-based features to form a hybrid feature set for predicting protein-protein interactions. A systematic comparison is performed between models trained on frequency features alone and those using a hybrid representation. The comparison showed that incorporating one-dimensional convolutional neural network-derived latent features improved the models’ performance of predicting protein-protein interactions. The dataset was split into training, validation, and test sets. Nested cross-validation was employed, with inner loops for hyperparameter tuning and outer loops for model selection. The random forest classifier achieved the best performance, with a mean receiver operating characteristic-area under curve of 0.91 and a test F1-score of 0.87. These results highlight the effectiveness of integrating deep feature learning with ensemble methods for predicting protein-protein interactions and build upon previous work focused on this fundamental problem.

**Author Summary:** Protein-protein interactions are fundamental in all biological processes. However, predicting these interactions is a key problem in molecular biology. Computational approaches have been tested to address this problem. We applied a mix of machine learning and deep learning to gain insight into the qualities of proteins that engage in interaction. First, we trained a deep learning model, which automatically learned the primary sequence and characters related thereto, reducing bias in the actual prediction process. We combined these features, or latent representations, with amino acid frequency features of protein sequences, and called the two together “hybrid features.” Then we performed a systematic comparison of amino acid frequency features-only with hybrid features, among four different machine learning classifiers. Our results suggest that the random forest classifier performed best among all four classifiers at predicting interactions between proteins. We propose that this approach could be used to improve efficiency in testing protein-protein interactions at the bench and may have applications to other biologically relevant molecular interactions.

## Introduction

Proteins perform various metabolic functions in living organisms and in doing so typically interact with other molecular components of the cell to enable diverse cellular and biological functions [1]. Many experimental methods have been developed to study protein-protein interactions (PPIs). Examples include the yeast two-hybrid system [2,3] and co-immunoprecipitation followed by mass spectrometry [4–6]. Experimental techniques have contributed to the generation of large databases containing datasets of PPI pairs. Notable examples are the Database of Interacting Proteins [7], the Mammalian Protein-Protein Interaction Database [8], the Biomolecular Interaction Network Database [9], the IntAct molecular interaction database [10] and the Molecular INTeraction database [11].

Experimental molecular techniques to detect PPIs are costly, in both time and money. In some cases, they are also subject to intrinsic bias resulting from experimental design. In contrast, computational approaches are relatively inexpensive and can be easily and dynamically adjusted to address concerns of classification bias [12,13]. In silico approaches to the prediction of protein-protein and protein-ligand binding are prominent in the field of bioinformatics [14]. From a validation standpoint, computational approaches can be used to filter the candidate pool of interacting proteins to high-confidence candidates, thus increasing experimental efficiency. In many cases, that increase in efficiency could, in principle, translate to greater depth of study. Several attempts to bioinformatically characterize PPIs using machine learning models have been made with varying degrees of success. For example, a 2014 study by Saha et al. demonstrated that gene ontology, coexpression and other metadata could be used along with an ensemble of a support vector machine, random forest (RF), decision tree (DT), and naive Bayesian classifiers to classify PPIs with high accuracy and high F1 values when applied to limited but balanced datasets [15]. While there was some utility in that method, it suffered from less than desirable F1 values (56%-73% for human and 20%-40% for yeast) when wider-spectrum, unbalanced, real-world data were used. This outcome was primarily the result of low recall and thus insufficient ability to correctly identify true positives; it deductively represented an overfit model [15,16].

Deep learning has also been used in PPI prediction. Soleymani et al. has provided a rigorous review on the use of deep learning to predict PPI [17]. A well-known deep learning method, termed DeepPPI used a deep neural network to one-hot classify PPIs. By plotting the features used in each hidden layer of the neural network, Du et al. were able to illuminate high-level discriminative features among various protein descriptors, including dipeptide frequency, amino acid composition and physicochemical descriptors [16]. While DeepPPI performed slightly less well than the ensemble method [15] at classifying a limited dataset, it performed similarly on extensive, imbalanced, real-world data (F1 = 71%-72% for human and 34%-36% for yeast). Additionally, it used basal level, sequence-based features rather than derived metadata. Therefore, the model has greater utility in that it can be applied to primary sequence data without extensive prior analyses. Another method employed an ordinal regression and recurrent convolutional neural network (RCNN) framework to predict PPIs based on interaction confidence scores [18]. The architecture comprised two RCNN encoders which shared the same parameters to extract robust local features and sequential information from protein pairs.

Following this extraction, one novel embedding vector was obtained by element-wise multiplication of the two embedding vectors from RCNNs. The second part of the architecture performed an ordinal regression model via multiple sub-classifiers that use the ordinal information behind the confidence score. Finally, PPIs were predicted by applying a threshold to the confidence score. This RCNN approach demonstrated improved performance in feature description compared with earlier models such as autocovariance [19] and composition transition distribution descriptor [20], and improved performance in predicting interactions compared to RF [21] and a support vector machine. However, the RCNN was built using bidirectional gated recurrent units, which suffer from slow convergence and low learning efficiency [22]. In many earlier studies, amino acid frequency-based features, and other engineered features were extracted and directly used to train machine learning classifiers [23–25]. However, constructing such features is time-consuming and requires substantial domain expertise. More recent approaches have explored deep learning for automatic feature extraction, often relying on recurrent neural network architectures.

While often overshadowed by recurrent or attention-based architectures, one-dimensional convolutional neural networks (1D CNNs) remain highly effective in sequence-based pattern recognition, due to their ability to automatically learn local spatial dependencies in sequence data [26,27], thereby reducing biases introduced by manual feature selection. Additionally, the representational quality of features learned through deep learning is rarely evaluated independently of downstream classification tasks. Poorly learned features may fail to capture meaningful biological signals [28], while excessively high-dimensional representations may introduce noise that obscures relevant patterns [29]. Therefore, both feature extraction and classification must be carefully optimized to achieve robust PPI prediction.

To address these challenges, we propose an integrated framework that employs a 1D CNN autoencoder for feature extraction, followed by systematic evaluation using multiple machine learning and deep learning classifiers. The 1D CNN autoencoder, in addition to extracting features from primary sequence data, evaluates the extracted features by reconstruction error during a pretraining phase; its ability to faithfully reconstruct primary sequence data indicates the characteristics it has learned are meaningful. Notably, these learned features are generalizable and can be applied to a range of tasks, including predicting PPIs, protein-ligand interaction, and broader drug discovery applications. In this study, however, we focus specifically on the prediction of PPIs. We further compare models trained using amino acid frequency alone with those utilizing a hybrid representation that combines amino acid frequencies with learned latent embeddings. We found that the incorporation of the hybrid representation substantially improved the models’ performance, and that the RF classifier outperformed other classification models. We propose that this integrated approach will lead to improved predictive performance.

## Methods

### Overview

This study employs a two-stage methodology to construct a robust PPI prediction framework, as shown in Fig 1. The first step utilizes a sequence-to-vector, 1D CNN autoencoder to learn compact latent representations of individual protein sequences directly from their primary amino acid sequences and the calculation of a 21-dimensional vector representing the frequency of the 20 biologically active amino acids and unknown residues. The second step involves training multiple classification models using different feature representations derived from step one. Briefly, models were trained using (i) amino acid frequency features alone and (ii) a hybrid feature set combining amino acid frequency features with latent representations derived from the 1D CNN autoencoder. This comparative analysis enabled systematic evaluation of the added predictive value of deep learning-based sequence representations over composition-based sequence descriptors alone.

**Fig 1.**
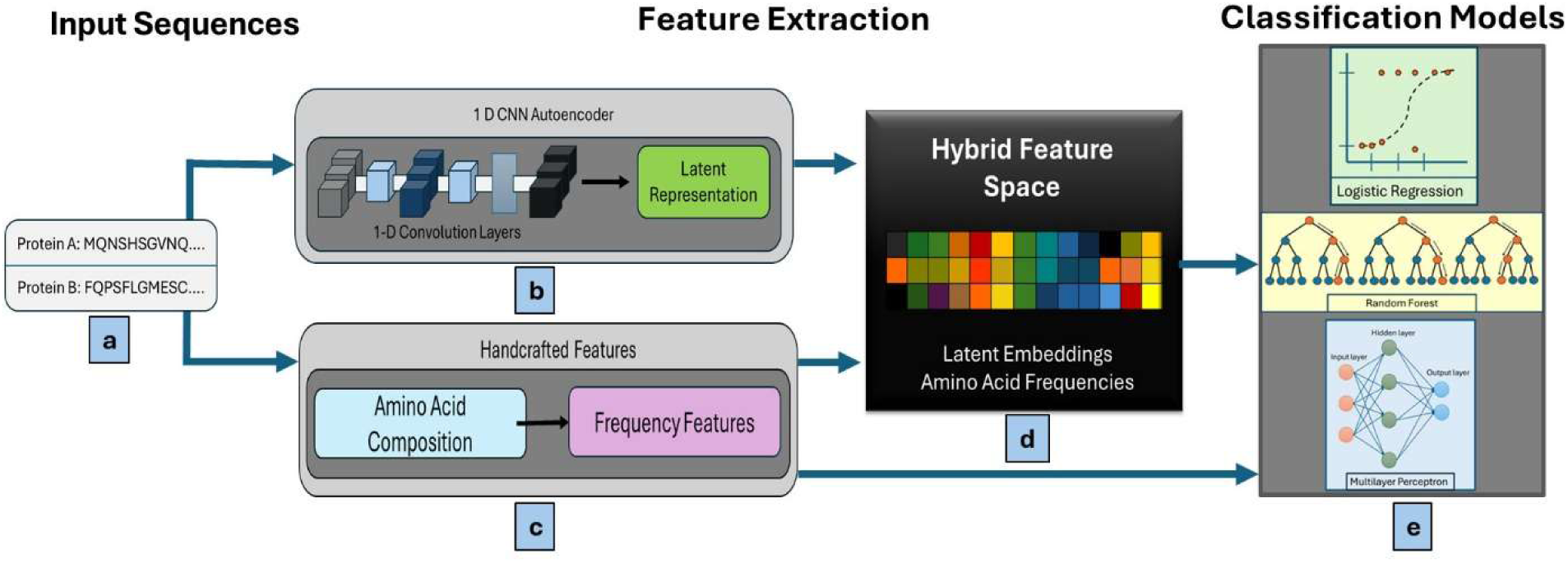
Schematic overview of the described sequence-based PPI prediction pipeline. (a) Primary amino acid sequences of two proteins are used as inputs. Feature extraction is performed through two parallel pathways: (b) a 1D CNN autoencoder that learns compact latent representations directly from protein sequences, and (c) frequency-based amino acid composition features that capture global sequence characteristics through frequency-based descriptors. (d) A hybrid feature space is constructed by the combination of latent embeddings generated by the 1D CNN autoencoder, concatenated with amino acid frequency features. (e) Different machine and deep learning classification models are independently trained using amino acid frequency features alone to enable direct performance comparison with hybrid representation. Both the hybrid feature space and the amino acid frequency-based feature set are evaluated using multiple classifiers, including logistic regression, random forest, and multilayer perceptron, to predict PPIs.

### Data collection and preprocessing

The first step when developing PPI prediction models is to select benchmark data for training, validation, and final testing. For the purpose of this study, we chose the *Saccharomyces cerevisiae* PPI data [16]. Du et al. (2017) preprocessed the original Database of Interacting Proteins data [7] by excluding records containing proteins with fewer than 50 amino acids and excluding homologs as identified by a CD-HIT clustering program with a threshold level of 40% identity, thus limiting the data to representative sequences and removing redundant sequences, which could disproportionately affect model performance metrics. Preprocessing resulted in a dataset reduction from 22,975 PPI pairs to 17,257. Noninteracting, or negative, protein pairs are difficult to define but essential for successful model training. One identification method is to infer that two proteins do not interact if they are not found in the same location within a cell [30]. To that end, Du et al. [16] extracted subcellular localization information for the proteins within their dataset from Swiss-Prot (http://www.expasy.org/sprot/). Proteins with non-overlapping subcellular localizations were assembled into non-interacting pairs. They generated a total of 48,594 negative pairs. From their data, we selected a random subset of 5,000 negative and 5,000 positive PPI pairs. Our dataset was limited to this size due to the substantial computational cost associated with developing a 1D CNN. Following random record selection, the dataset was split into training, validation, and test sets using a 70:15:15 ratio. The test set was kept isolated from all training and validation data, thus ensuring that final model evaluation could be performed on unseen data.

### Feature extraction

Each of the 20 standard amino acids in a protein sequence was encoded using a unique integer from 1 to 20, with an additional index (21) used for ambiguous or unknown residues. This encoding resulted in conversion of each protein sequence into an x-dimensional vector, where x represents the length of the protein sequence. To obtain uniform input dimensions for neural network training, all sequences were then padded to the length of the longest sequence in the dataset. The fully encoded/padded sequences were then utilized to train a 1D CNN autoencoder enabling the model to capture features representing the spatial relationships among sequential amino acids. The encoder comprises a 32-dimensional embedding layer, followed by a 1D convolutional layer with 64 filters (kernel size = 3) and rectified linear unit [31] activation and a global max-pooling layer [32]; the resulting representation is then passed through a fully connected layer to obtain a 400-dimensional latent vector. The decoder then reconstructs the input sequence using a RepeatVector layer, a Long Short-Term Memory layer with 64 units, and a TimeDistributed dense layer with softmax activation over the 21-token vocabulary.

The autoencoder was trained using an Adaptive Moment Estimation optimizer with categorical cross-entropy loss for up to 50 epochs (batch size = 128), employing early stopping based on training/validation loss to detect overfitting. This configuration ensures that the learned latent vectors capture informative sequence patterns while maintaining stable generalization performance. Training dynamics were assessed using learning curves comparing training and validation loss and accuracy across epochs, providing insight into model convergence and generalization behavior.

In addition to feature extraction using the 1D CNN, frequency-based features were derived from the counts of each of the 20 standard amino acids in each protein sequence. This procedure was applied to each protein that comprises negative and positive PPI pairs. Therefore, the frequency-based features for each PPI pair consist of two 20-dimensional amino acid composition vectors that characterize the frequency distribution of residues in each protein.

### Machine learning

Four algorithms - three machine learning and one deep learning - were evaluated using the aforementioned feature sets. These included logistic regression (LR), DT, RF and multi-layer perceptron (MLP). Each model was evaluated in terms of accuracy, precision, recall [33,34]. In all cases, F1-score was considered the primary evaluation metric for model quality. In addition to these primary performance metrics, ROC-AUC was calculated for each model. The equations used for performance metrics and AUC calculations are contained in Equations 1-7. Each model was 3-fold cross-validated during hyperparameter tuning, resulting in mean performance metrics. A 3-fold cross-validation strategy was selected to balance computational efficiency with robust model evaluation across all classifiers. In our framework, hyperparameter optimization requires repeated training of multiple models (LR, DT, RF, and MLP) over high-dimensional feature representations derived from both amino acid frequency features and 1D CNN-based embeddings. Increasing the number of folds (e.g., 5- or 10-fold cross-validation) would substantially increase computational cost due to repeated model fitting across numerous hyperparameter configurations. Given the use of stratified splitting and consistent performance across folds, 3-fold cross-validation provides a reliable and computationally efficient estimate of model performance. The optimal model was then evaluated using the test set data.

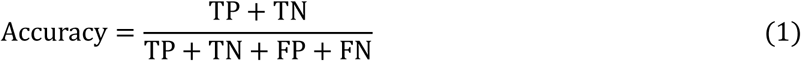

Where TP, TN, FP, and FN represent true positive, true negative, false positive, and false negative, respectively.

Precision measures the proportion of true positive predictions out of the total positive predictions and is defined in Equation 2,

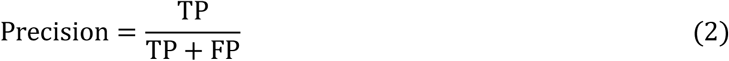

Recall calculates the proportion of correctly predicted positive instances out of all actual positive instances, given by Equation 3,

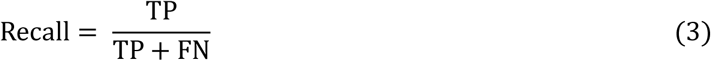

The F1 score, which is the harmonic mean of precision and recall, is calculated as follows in Equation 4,

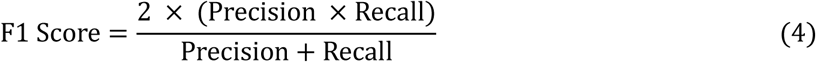

In addition, ROC-AUC is used to assess the model’s ability to discriminate between classes across all possible classification thresholds. The ROC curve plots the true positive rate (TPR) against the false positive rate (FPR), where,

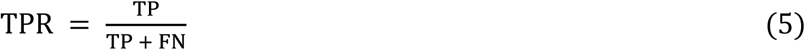

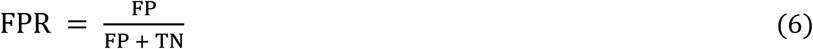

The ROC-AUC is defined as the area under this ROC curve and yields a threshold-independent measure of separability:

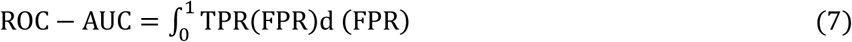

ROC-AUC ranges from 0 to 1, where a value of 0.5 indicates performance comparable to random guessing, and values closer to 1.0 indicate stronger class separability [35,36]. Unlike accuracy, which depends on a single decision threshold, ROC-AUC summarizes the ranking quality of predicted probabilities/scores and is especially informative when comparing models under varying threshold choices or in settings where the operating point may change [37,38].

LR was used as a baseline classifier to assess the model’s capability for PPI prediction. As a linear probabilistic model, LR estimates the likelihood of interaction as a sigmoid transformation of a weighted linear combination of input features, providing a well-established and interpretable benchmark for evaluating more complex nonlinear models. Because LR is sensitive to feature scaling [39], input features were standardized prior to model fitting to ensure numerical stability and balanced coefficient estimation.

The model was implemented using a scikit-learn [40] pipeline consisting of z-score normalization via StandardScaler [40] followed by an LR classifier with a maximum of 1,000 training iterations. Hyperparameter optimization was performed using an exhaustive grid search with 3-fold cross-validation. The inverse regularization strength was varied across five values, C∈{0.01,0.1,1,10,100}, allowing exploration of both strongly and weakly regularized models. To ensure a stable convex optimization setting, the L2 penalty and the limited-memory Broyden-Fletcher-Goldfarb-Shanno (LBFGS) solver were used for all configurations [40]. Model selection was conducted using a multi-metric evaluation strategy in which both F1-score and accuracy were computed for each fold; however, the optimal hyperparameter configuration was selected by maximizing the mean cross-validated F1-score, reflecting the importance of balancing precision and recall under class imbalance. The best-performing model was subsequently retained for final evaluation.

A DT classifier was employed to capture non-linear interactions that cannot be represented by linear classifiers [41]. DTs recursively partition the feature space by selecting split points that maximize class purity, enabling the model to learn nonlinear decision boundaries and interaction effects among input variables [42]. DT hyperparameter optimization was conducted using Randomized Search cross-validation (CV) with stratified 3-fold cross-validation. The search space was defined over key structural parameters controlling model complexity and generalization. Specifically, the maximum tree depth was sampled from a discrete uniform distribution of range 1 to 10 while both the minimum number of samples required to split an internal node and the minimum number of samples required at a leaf node were sampled from {5,…,9}. The optimal model was selected by maximizing the mean cross-validated F1-score, reflecting the importance of balanced precision and recall. The resulting best-performing decision tree was retained for subsequent evaluation. RF classifiers construct an ensemble of decision trees trained on bootstrap samples of the data, while additionally decorrelating trees by selecting a random subset of features at each split. Final predictions are obtained by aggregating the outputs of individual trees via majority-rule voting, which generally improves generalization performance in high-dimensional feature spaces such as PPI representations [43].

For RF hyperparameter optimization, a Randomized Search CV with stratified 3-fold cross-validation and a multi-metric evaluation strategy was used. Trees were randomly sampled from a discrete uniform distribution; n_estimators ∈ {5,…,14}, and the maximum tree depth was sampled from a range of 1 to 4. To control node-level regularization, both the minimum number of samples required to split an internal node and the minimum number of samples required in a leaf node were sampled from {5,…,9}. Feature subsampling at each split was tuned using max_features ∈{sqrt, log, None). A total of 30 randomly sampled hyperparameter configurations were evaluated. For each configuration, both F1-score and accuracy were computed; however, the optimal model was selected by maximizing the mean cross-validated F1-score (refit=“f1”). The best-performing RF was retained for subsequent evaluation.

An MLP classifier was used to evaluate the performance of a fully nonlinear neural model on the amino acid frequency-based PPI features. Unlike tree-based methods, the MLP learns distributed feature representations through stacked, fully connected layers [44], enabling it to capture high-order nonlinear interactions among input variables [45]. Because neural networks are sensitive to feature scaling [46], all input features were standardized prior to model fitting.

The MLP was implemented using a scikit-learn pipeline consisting of z-score normalization followed by a feedforward neural network trained with the Adaptive Moment Estimation optimizer. Hyperparameter optimization was performed using Randomized Search CV with stratified 3-fold cross-validation and a multi-metric evaluation strategy. Network architecture was explored by sampling hidden-layer configurations (16,8), (8,4), (4,2), corresponding to two-layer networks of decreasing capacity. The L2 regularization strength was sampled from a log-uniform distribution in the range (10^-4^,1), and the initial learning rate was sampled from η∈ (10^-4^,10^-2^). The rectified linear unit activation function [31] was used for all configurations. To mitigate overfitting and ensure stable convergence, early stopping was enabled with a validation fraction of 0.15, and training was terminated if no statistically significant improvement was observed for 10 consecutive iterations. A total of 30 random hyperparameter configurations were evaluated, and the optimal model was selected by maximizing the mean cross-validated F1-score; accuracy was also recorded for reference.

## Results

The performance of a 1D CNN autoencoder was evaluated in terms of its reconstruction error to verify its efficacy in learning informative latent representations of higher dimensional protein sequence data. Performance metrics are presented for LR, DT, RF and MLP classifiers, trained on both the 1D CNN-derived features alone and those same features combined with amino acid frequency features.

### 1D CNN training performance

The 1D CNN autoencoder (Fig 2) was trained for 35 epochs. Training and validation loss were sharply reduced during the first ∼5 epochs, indicating that the risk of misclassification rapidly decreased due to parameter optimization. After this phase, both loss curves flattened and remained near-constant, suggesting the optimizer was making small-scale adjustments that resulted in modest improvement (Fig 3). The complementary pattern between training and validation accuracy increased quickly and then stabilized around ∼0.88–0.89. The small and stable generalization gap throughout training suggested the model’s capacity and regularization were sufficient to minimize fitting noise in the training set. In other words, there was no systematic divergence between training and validation metrics that would indicate overfitting.

**Fig 2.**
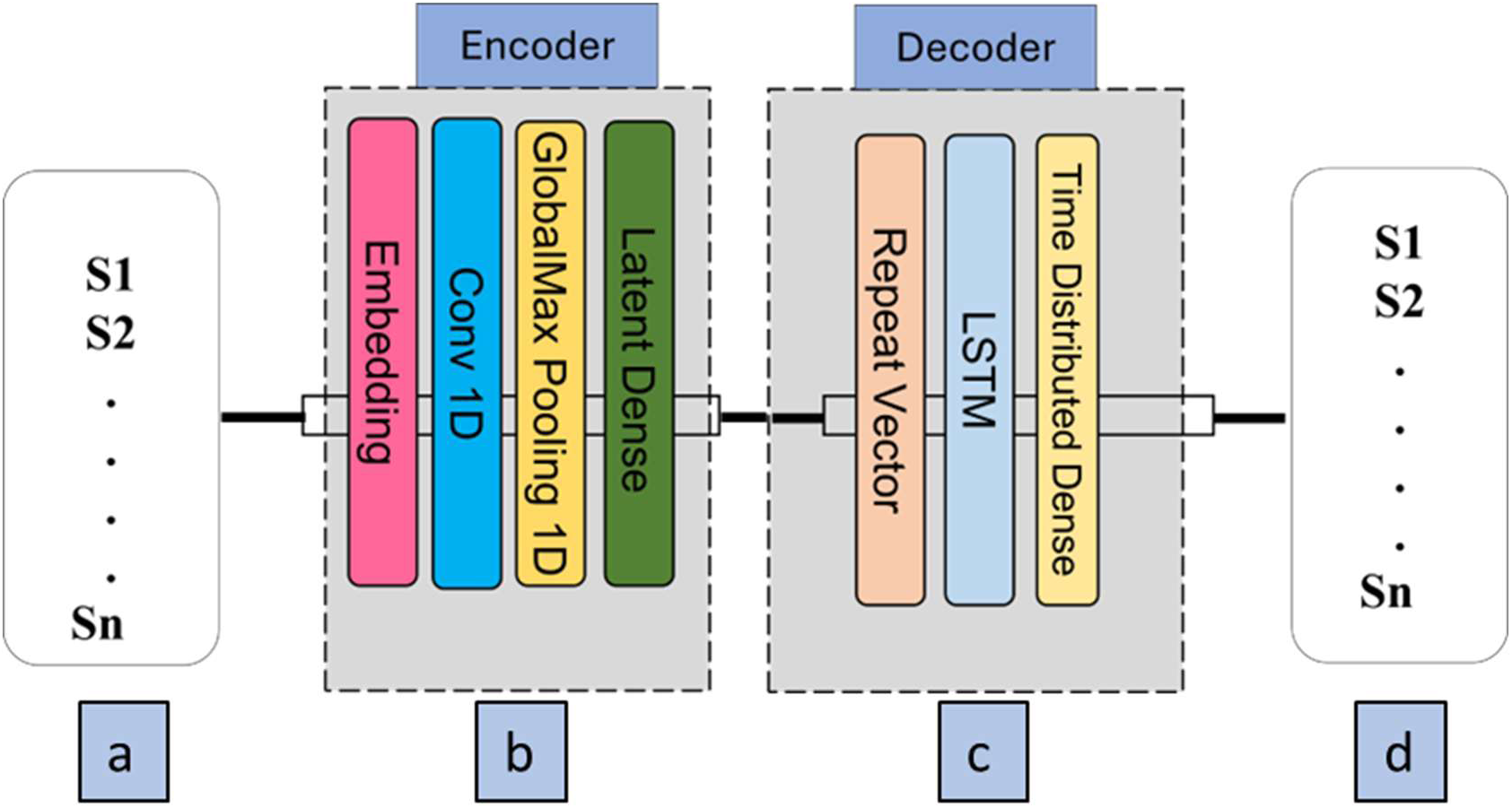
Architecture of the 1D CNN-Long Short-Term Memory autoencoder used for protein sequence representation learning. (a) Input protein sequences (S1-Sn) were provided as raw amino acid sequences. (b) Encoder module: sequences were first transformed through an embedding layer, followed by a one-dimensional convolutional (Conv 1D) layer to capture local sequence motifs. GlobalMax Pooling 1D reduced dimensionality while preserving salient features, and a dense latent layer generated a compact sequence representation. (c) Decoder module: the latent vector was repeated and passed through a Long Short-Term Memory layer to model sequential dependencies, followed by a time-distributed dense layer to reconstruct the original sequence. (d) Output reconstructed sequences (S1-Sn) enabled unsupervised training via reconstruction loss. The encoder-derived latent representations were subsequently used as learned sequence features for downstream PPI prediction.

**Fig 3:**
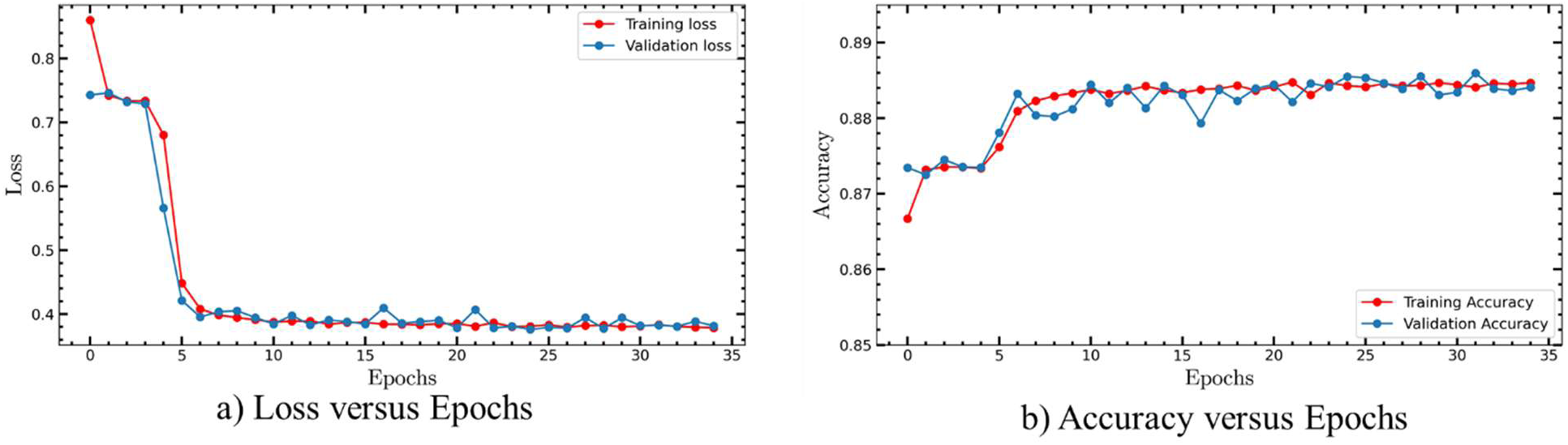
Learning curves for stage 1 of the framework: 1D CNN autoencoder. (a) Both training and validation loss decreased rapidly within the first five epochs and subsequently stabilized at a low value, indicating fast convergence of the network. (b) Training and validation accuracy increased sharply during the initial seven epochs and remained consistently high.

### Machine learning

To quantify the contribution of the 1D CNN embeddings, models were optimized using 3-fold cross-validation during hyperparameter tuning. Identical model selection criteria were used across all feature sets. This comparison isolated the effect of feature representation while holding the learning algorithm and tuning protocol constant. Using amino acid frequency features alone, the optimized LR models achieved a mean inner-cross validated F1-score of 0.63. When LR models were trained using the hybrid feature representation, which combined amino acid frequency with 1D CNN-derived latent embeddings, an improved inner cross-validation performance, relative to the frequency only baseline, was achieved, with an F1-score of 0.66.

Throughout both trainings, the same hyperparameter grid and selection procedure were applied to ensure consistency. For the frequency only feature representation, the optimized DT achieved a mean inner cross-validated F1-score of 0.66. When DT was trained using the hybrid feature representation, an improved inner cross-validation performance was observed. The best-performing configuration for DT employed a maximum tree depth of 9, a minimum of 7 samples required to split an internal node, and a minimum of 5 samples per leaf. Under inner three-fold cross-validation, this configuration achieved a mean F1-score of 0.69. The optimized RF classifier achieved a mean inner cross-validated F1-score of 0.76. Hyperparameters were selected using randomized search over ensemble size, tree depth, node regularization parameters, and feature subsampling strategies, with model selection based on mean F1-score. When trained with the same hyperparameters using the hybrid feature representation, RF achieved substantially improved inner cross-validation performance, with a mean F1-score of 0.81. For the frequency-only feature set, the optimized MLP achieved a mean inner cross-validated F1-score of 0.66. The best-performing configuration employed two hidden layers with 16, 8 neurons, an L2 regularization strength of α = 0.0031, and an initial learning rate of 0.00023, selected via randomized search over network architecture, regularization strength, and learning rate. When trained using the hybrid feature representation, the MLP again selected a two-layer architecture with 16, 8 neurons, but with different optimal regularization strength and learning rate. The best-performing configuration employed α = 0.00053 and an initial learning rate of 0.0017, yielding improved inner cross-validation performance relative to the frequency-only baseline, F1-score 0.69. This shift in optimal regularization was attributed to the increased representational capacity introduced by the CNN-derived latent features (Table 1).

**Table 1:**
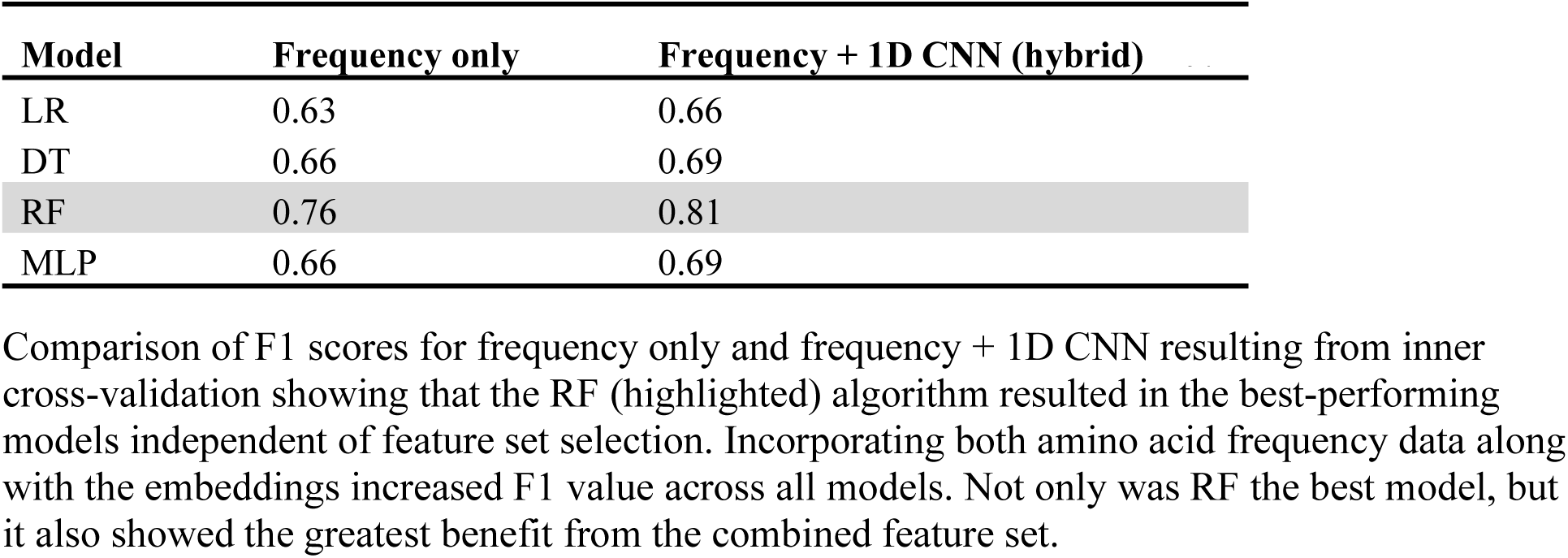
Comparison of F1 scores among different classifiers using inner 3-fold CV.

### Feature selection

Across all evaluated classifiers, the inclusion of 1D CNN–derived latent features led to consistent improvements in inner cross-validated performance compared to amino acid frequency features alone. The magnitude of improvement was modest for linear models, more pronounced for nonlinear classifiers, and largest for ensemble-based methods. These findings confirmed that learned sequence representations capture interaction-relevant information that is not fully encoded by amino acid composition, justifying their inclusion in classification models.

### Comparative evaluation of models using frequency + 1D CNN features

Inner cross-validation results consistently demonstrated that incorporating 1D CNN–derived latent features alongside amino acid frequency features led to improved predictive performance across all evaluated classifiers compared to frequency-only representations. These improvements were observed for linear, nonlinear, and ensemble-based models, indicating that the learned CNN embeddings captured interaction-relevant sequence information not fully represented by amino acid composition alone. Based on these findings, subsequent model comparisons and evaluations were conducted exclusively using the hybrid, frequency + 1D CNN feature representation, which was selected as the most informative input space for downstream analysis.

LR, DT, RF, and MLP classifiers were evaluated using outer 3-fold stratified cross-validation under balanced class partitions and consistent evaluation criteria. For each model, mean and standard deviation of training and validation ROC–AUC were computed across folds to assess both discriminative performance and generalization stability (Table 2).

**Table 2:** Outer 3-Fold Cross-Validation Performance Using Frequency + 1D CNN Features.

| Model | CV Train ROC-AUC | CV Val ROC-AUC |
| --- | --- | --- |
| LR | $0.7723 \pm 0.0021$ | $0.7387 \pm 0.0062$ |
| DT | $0.8555 \pm 0.0166$ | $0.7865 \pm 0.0147$ |
| RF | $0.9899 \pm 0.0009$ | $0.9160 \pm 0.0064$ |
| MLP | $0.8889 \pm 0.0128$ | $0.7793 \pm 0.0103$ |

Performance is reported as mean ± standard deviation of ROC-AUC across folds for both training and validation sets. RF shows the best validation performance.

Classifiers capable of modeling nonlinear relationships exhibited superior validation performance compared to linear classifiers. RF achieved the highest validation accuracy and ROC–AUC, distantly followed by the MLP. LR and DT classifiers demonstrated comparatively low performance. These results indicate that the combination of 1D CNN embeddings with ensemble-based learning was especially effective for PPI prediction.

### Final model training and evaluation

Following initial model optimization, the RF classifier was selected as the algorithm to produce the final model. This selection was supported not only by its superior validation metrics but also by its stability across folds, indicating that the model’s performance was not driven by a small subset of stochastically favorable splits. The best hyperparameters that were identified during optimization were used to train the final RF model on a combined training and validation dataset, thus maximizing the amount of training data. On the combined training data, the model achieved an accuracy of 0.95, suggesting that the classifier effectively learned discriminative interaction patterns from the proposed hybrid feature representation and that neither false positives nor false negatives were noteworthy. The frequency-based features captured global compositional tendencies and coarse sequence statistics, whereas the 1D CNN embeddings encoded local motif-like patterns as well as higher-order dependencies.

When evaluated on the independent held-out test set, the model achieved F1-score of 0.87. The reduction from development to test performance was expected, as the combined training–validation results were obtained on data that directly contributed to fitting the final model, while the test set provided a stricter estimate of generalization to unseen protein pairs. Importantly, test performance remained high and consistent with the cross-validation trends.

To better understand the types of errors made by the classifier, a confusion matrix was constructed based on the test set (Table 3). Out of 750 true interaction pairs, 704 were correctly predicted as interactions, while 46 were misclassified as non-interactions (false negatives). Out of 750 true non-interactions, 611 were correctly predicted as non-interactions, while 139 were incorrectly predicted as interactions (false positives). This pattern indicated that the model was slightly more prone to predicting interactions for some non-interacting pairs than missing true interactions.

**Table 3:**
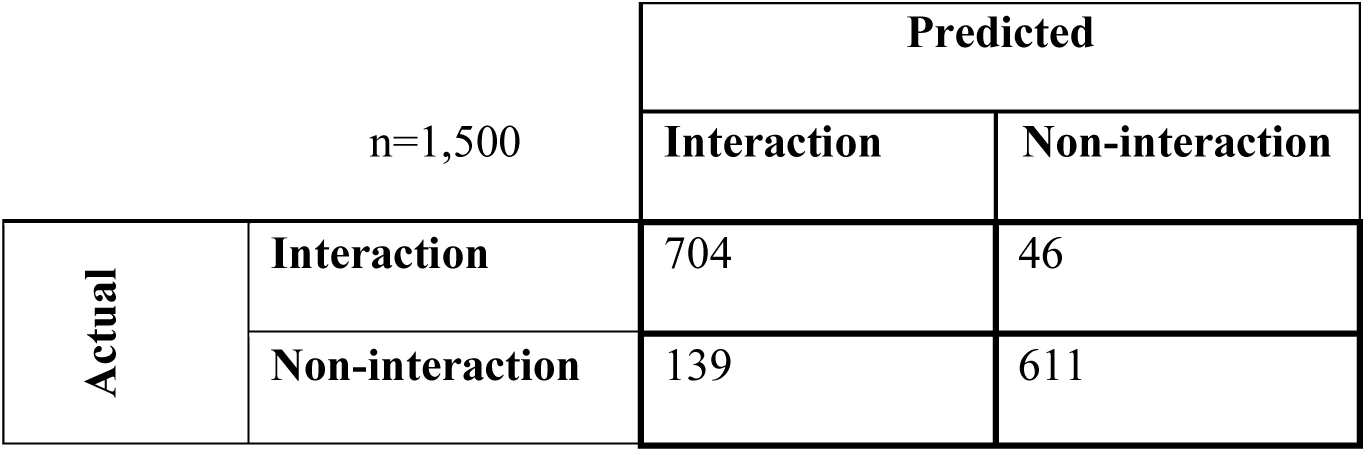
Confusion matrix of the final model on the independent test set.

Class-specific precision, recall, and F1-scores were calculated for the test set (Table 4). For the interaction class, the model achieved a precision score of 0.83, recall of 0.93, and F1 of 0.88, indicating strong sensitivity (i.e., most true interactions are recovered) with a moderate number of false positives. For the non-interaction class, the model achieved precision of 0.93, recall of 0.81, and F1 of 0.86, showing that when the model predicted a non-interaction it was usually correct, although some non-interacting pairs were still classified as interactions. Taken together, the confusion matrix and per-class metrics suggested a favorable trade-off: the classifier prioritized capturing true interactions (high recall for interactions) while maintaining strong overall discrimination between the two classes.

**Table 4:** Class-wise precision, recall, and F1-score of the final model on the independent test set.

| Class | Precision | Recall | F1-score |
| --- | --- | --- | --- |
| Interaction | 0.83 | 0.93 | 0.88 |
| Non-interaction | 0.93 | 0.81 | 0.86 |

Evaluation metrics for both classes on the test set showing that precision was higher for the negative PPI class, recall was higher for the positive PPI class and overall, the combination of precision and recall in the form of F1-score indicates that the model was slightly better at classifying the positive PPI class. The better metrics are shaded.

## Discussion

In this study, an integrated sequence-based framework for PPI prediction was developed by combining deep representation learning with four machine learning algorithms (LR, DT, MLP, and RF). A 1D CNN autoencoder was used to create compact latent representations (embeddings) of the amino acid sequences to capture patterns within the sequences. Those embeddings were then concatenated with amino acid frequency features to create a hybrid feature space. This strategy enabled the simultaneous exploitation of local patterns in the sequences and global composition characteristics. Our results demonstrate that the inclusion of features derived from a 1D CNN consistently improved predictive performance across all examined classifiers when compared to amino acid frequency features alone. The observed performance gains were more apparent for nonlinear classifiers and most pronounced for ensemble methods.

Based on the superior cross-validation performance of the hybrid feature representation, we focused exclusively on models trained using the combined frequency and 1D CNN features. The RF classifier achieved the highest validation accuracy and ROC–AUC among all evaluated models, demonstrating both strong discriminative capability and stable generalization. RF is well-suited to this setting because it models nonlinear interactions between heterogeneous feature groups, is comparatively robust to irrelevant/weak predictors [47], and typically requires limited feature scaling [48] or distributional assumptions [43]. Thus, the observed strong development-set performance indicates that the learned representation provides a separable structure for the interaction vs. non-interaction classes.

The final RF model, trained on the combined training and validation datasets from cross validation, further exhibited robust performance on test data, indicating effective learning of interaction patterns from the hybrid feature space. These results collectively demonstrate that utilization of 1D CNN sequence-based embeddings and an RF classifier improved the PPI predictive ability relative to other reported methods in terms of accuracy, F1 score and ROC-AUC. Shen et al. [49] built a support vector machine classifier based on physicochemical properties that achieved a mean accuracy of 83.90% (F1 = ∼85%) across all test sets. Our RF model exceeded those metrics with an accuracy of 88.09% (F1 = ∼86.50%), further demonstrating the utility of using sequence-based 1D CNN embeddings along with amino acid frequency data. An added benefit is that our method obviates the need for the physicochemical encoding used in the Shen et al. method and results in greater classification confidence for downstream applications.

Both MLP and RF performed well. RF resulted in the best performance, in terms of ROC-AUC. The capacity of RF to achieve the highest ROC-AUC, and its consistent performance across folds, indicates that our hybrid feature space, consisting of both amino acid frequency data and a sequence-based 1D CNN embeddings exposes complex nonlinear interactions between individual features that were leveraged for RF classification [43]. Ensemble models such as RF are better at capturing nonlinear interactions than other models. Moreover, they minimize issues associated with small sample size by averaging the signal across sub-samples and multiple paths through the model. This minimizes the potential for overfitting [50] and thus further increases confidence in predicting PPI. We propose that, from a practical standpoint, our RF model can be used to objectively filter voluminous lists of potential protein partners to identify with high confidence likely PPIs, and that using this way will replace time-consuming experimental assays.

The confusion matrix for our RF model (Table 3) indicates that the model produced more false positives (139) than false negatives (46), suggesting a bias toward predicting interactions. This pattern is consistent with the class-wise evaluation metrics. Recall is higher for the interaction class (0.93), reflecting strong sensitivity, while precision is higher for the non-interaction class (0.93), indicating reliable identification of non-interacting pairs. Together, these results suggest a trade-off in which the model prioritizes the detection of true interactions. Such bias may be advantageous to an investigator when planning targeted studies, since failing to identify true interactions can be more detrimental than tolerating false positives [51]. To minimize the effects of false positives, a large number of true negatives in the training data is necessary. Unfortunately, validated true negatives are not easily obtained because the focus of the scientific community is generally on identifying interactions, not on identifying non-interacting pairs. Additionally, interactions are conditional. For example, interactions between viral coat proteins and integral membrane proteins on the surface of animal cells depend on their being exposed to each other [52]. The dataset used herein [16] includes positive interaction pairs that are derived from experimentally validated data, whereas the negative pairs included were inferred based on lack of cellular colocalization. This inference is likely to be generally accurate and appropriate for inferential purposes. However, there is also the possibility of stochastic or conditional interactions between proteins that are not normally colocalized - even within the same organism - which cannot be identified using a cellular colocalization criterion. Machine learning models are typically built under the assumption of ground truth within the input labels. That assumption may not fully hold for the negative class in this context and likely resulted in the elevated number of false positives. In other words, some of the false positives (Table 3) may actually be true positives despite lacking colocalization.

## Conclusion

The results of this study highlight the effectiveness of integrating deep sequence-level feature extraction with ensemble learning for PPI prediction. By leveraging the complementary strengths of learned latent embeddings and biologically interpretable frequency-based features, the proposed framework provides a scalable and annotation-independent approach for sequence-based PPI prediction. Future work will include the evaluation of attention-based architectures to better capture long-range dependencies, evaluate transferability across species and interaction databases, and extend the framework to multi-class or confidence-based interaction prediction tasks.

Beyond PPI, 1D CNN embeddings may be useful in capturing conserved motifs and functional regions in biological sequences. This would support tasks such as predicting binding sites, and identifying conserved regions associated with pathogenicity. One area where there is overlap of PPI and pathogenicity that warrants future investigation is the binding of surface-exposed viral proteins to their host receptors.

## Code and data availability

The code used in this study is available on Github.

## Acknowledgments

We would like to thank Dr. Anandi Dutta for her comments in the manuscript.

## Notes

### Competing Interest Statement

The authors have declared no competing interest.

